# Increased drug-seeking and vulnerability to relapse after escalation of nicotine intake in male and female rats

**DOI:** 10.1101/2025.09.14.676091

**Authors:** Kévin Letort, Laetitia Lageyre, Serge H. Ahmed, Karine Guillem

## Abstract

Nicotine addiction is characterized by escalated drug use, craving and a high relapse rate after abstinence. However, because of difficulties in demonstrating escalation of nicotine use in rats, its relation to other addiction-related phenomena is currently unknown. Recently, we showed that, compared to rats with a fixed moderate dose of nicotine, rats with access to increasing high doses of nicotine for self-administration progressively escalated their nicotine intake. Whether these animals with escalating patterns of nicotine self-administration also develop other behavioral signs of addiction remains to be investigated. Here we report that after escalation of nicotine intake, animals have a greater difficulty of abstaining from seeking the drug, a greater responsiveness to nicotine-induced craving-like behavior, and an increased vulnerability to re-escalate nicotine intake post-extinction than rats with stable patterns of nicotine intake. No substantial sex differences in the development of these different addiction-related phenomena were observed. Finally, after escalation, nicotine intake also became primarily dependent on nicotine reinforcement and less so on the nicotine-paired cue. Overall, this study shows that most of the post-escalation behavioral changes previously seen with other drugs of abuse are generalizable to nicotine intake escalation.

## Introduction

With 1.3 billion tobacco users worldwide, nicotine addiction remains the most prevalent addiction in the world today and a leading cause of preventable morbidity and mortality [1]. Like other addictions, nicotine addiction is characterized by escalated drug use, craving and a high relapse rate after abstinence. However, unless subjected to different access conditions [2,3], nicotine intake escalation has been difficult to establish in rodents. Interestingly, we have recently developed a robust rat model of escalation of nicotine intake using a differential dose procedure that produced two patterns of nicotine intake in rats [4]. Specifically, with access to a fixed, moderate unit dose of nicotine for self-administration (i.e., 30 μg/kg/injection), drug intake remained low and stable while, in contrast, with access to an increasingly large dose of nicotine for self-administration (i.e., from 30 μg to 240 μg/kg/injection), drug intake gradually escalated over days.

Importantly, previous research with other drugs (i.e., cocaine, heroin or methamphetamine) has consistently shown that rats with escalating patterns of drug self-administration also develop other addiction-related behavioral alterations that are not observed in controls animals with stable pattern of drug use [5–9]. In particular, rats that have escalated their drug intake i) have a greater difficulty of abstaining from seeking the drug, as measured by increased responding on the drug-paired lever during extinction, ii) are more responsive to nicotine-induced craving-like behavior, as measured during reinstatement of drug seeking after extinction, and iii) have an increased vulnerability to relapse, as shown a rapid re-escalation of drug intake when access to the drug is made again available. Whether these behavioral alterations are also observed after nicotine intake escalation remains to be investigated.

Here, we used the differential dose procedure [4] to directly compare the effects of stable versus escalation pattern of nicotine self-administration on several nicotine-taking and - seeking behaviors relevant to addiction. Specifically, we compared levels of drug intake, drug-seeking behavior during extinction, nicotine-induced reinstatement of nicotine-seeking behavior, and resumption of nicotine intake after extinction in both male and female rats. Moreover, as response contingent cues accompanying nicotine taking are known to influence nicotine self-administration [10–13], we also examined the contribution of nicotine and nicotine associated-cues in maintaining nicotine-taking and -seeking behaviors after nicotine intake escalation.

## Material and Methods

### Subjects

A total of 41 adult male and 28 female Wistar rats were used (225–250g for the males and 200-220g for the females at the beginning of experiments, Charles River, Lyon, France). 12 male and 5 female rats did not complete the entire experiment because of loss of catheter patency, thereby leaving 29 male and 23 female rats for the final analysis. Rats were housed in groups of 2 and were maintained in a light-(reverse light-dark cycle), humidity-(60 ± 20%) and temperature-controlled vivarium (21 ± 2 °C), with water and food available ad libitum. All behavioral testing occurred during the dark phase of the light-dark cycle. Home cages were enriched with a nylon gnawing bone and a cardboard tunnel (Plexx BV, The Netherlands). All experiments were carried out in accordance with institutional and international standards of care and use of laboratory animals [UK Animals (Scientific Procedures) Act, 1986; and associated guidelines; the European Communities Council Directive (2010/63/UE, 22 September 2010) and the French Directives concerning the use of laboratory animals (décret 2013–118, 1 February 2013)]. The animal facility has been approved by the Committee of the Veterinary Services Gironde, agreement number A33-063-952.

### Apparatus

Fourteen identical operant chambers (30 × 40 × 36 cm) were used for nicotine self-administration testing and training (Imetronic, Pessac, France). They were individually enclosed in wooden cubicles equipped with a white noise speaker (45 ± 6.2 dB) for sound attenuation. Each chamber was equipped with two retractable metal levers on opposite panels of the chamber and a corresponding white cue light above each lever. Each lever was connected to a syringe pump placed outside, on the top of the cubicle. Each self-administration chamber was also equipped with two pairs of infrared beams above the grid floor. Both pairs crossed the chamber on its length axis allowing to count the number of horizontal displacements (crossovers). Operant chambers were connected to a PC via an Imetronic interface and experiments were controlled and conceived using an Imetronic software (Imetronic, France).

### Surgery

Under deep anesthesia (intraperitoneal injection of a mixture of ketamine, 100 mg/kg and xylazine, 15 mg/kg), rats were surgically implanted with a silastic catheter (Dow Corning Corporation, Michigan, USA) in the right jugular vein. The catheter was tunneled subcutaneously and exited approximately 2 cm below the scapulae at the midline of the back. To ensure catheter patency, rats received a daily flush of 0.2 ml of an ampicillin solution (0.1 g/ml) containing heparin (300 IU/ml) post-surgery. After the surgery, animals were weighed every 2-3 days. Behavioral testing began 7-10 days after surgery.

### Experimental procedures

An overview of experimental designs is provided in Figure 1a.

**Figure 1.**
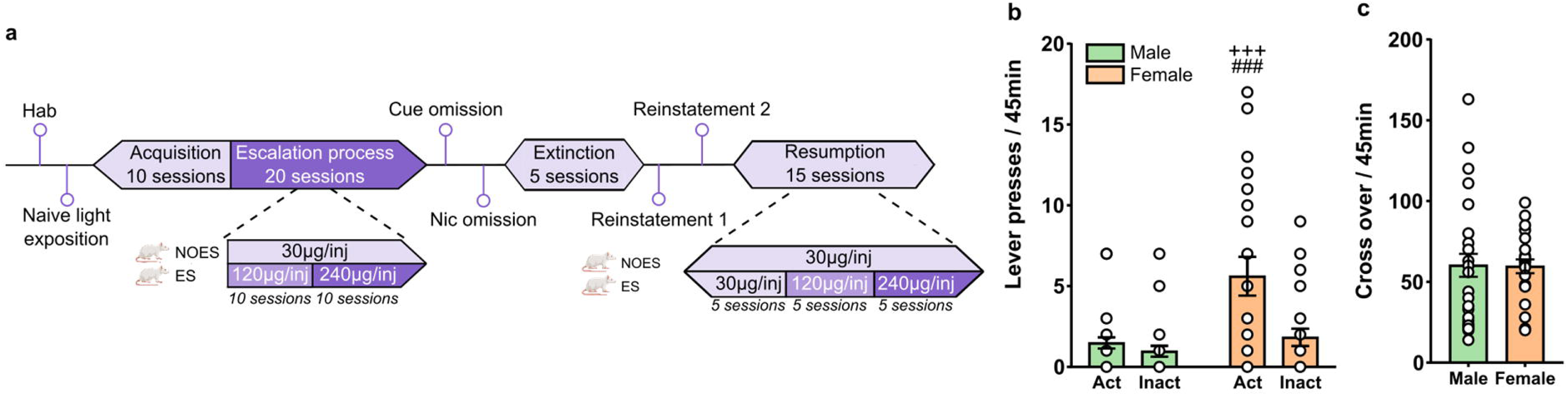
**(a) Sequence of experimental events.** Animals (n = 52) were first habituated (Hab) to the operant cages then completed one operant light self-administration session to measure the initial reinforcing value of the 20-s light cue (Cue exposure). Rats were next trained to press a lever to self-administer a unit dose of nicotine (30 µg/kg/injection free base; 10 days), then subjected to a differential dose procedure (20 days): animals in the no-escalation group (NOES, n = 26) had access to 30 μg/kg/injection for self-administration, while animals in the escalation group (ES, n = 26) had access to an increasing dose of intravenous nicotine (120 μg and 240 μg/kg/injection). Animals next underwent a cue omission (CueOm) and a nicotine reward omission session (NicOm) test session. They were then subjected to 5 days of extinction before nicotine-induced reinstatement of drug-seeking was assessed on two within-session reinstatement tests with increasing doses of nicotine (reinstatement 1 and reinstatement 2). Finally, rats were given again access to nicotine self-administration and escalation as before extinction for 15 additional days. **(b, c) Initial reinforcing value of the light cue in naïve rats.** (**b**) Mean (±SEM) number of lever presses on the active (*colored bars*) and inactive (*white bars*) lever, and (**c**) mean (±SEM) number of cross over during the light cue self-administration session in male (green) and female (orange) rats. ^+++^p<0.001, different from males; ^###^p<0.001, from the inactive lever.

#### Initial reinforcing value of the light cue

Animals were first habituated to the operant cages for one 2h session. On the subsequent day, they completed one 90-min operant light self-administration session to measure the initial reinforcing value, if any, of the 20-s light cue to be associated with nicotine in naïve animals. The session started with the extension of two levers and consisted of two periods of 45 min each. During the first 45-min period, responding on either lever had no programmed consequence. The onset of the second 45-min period was signaled by turning on the light-cue above one of the two levers for 20 s. Then, responding on this lever triggered this 20-s light-cue under a fixed-ratio 1 schedule; responding on the other lever had no scheduled consequence. Note that the lever paired with the 20-s light-cue will be the active lever during nicotine self-administration sessions (see below).

#### Acquisition and escalation of intravenous nicotine self-administration

All animals were first trained to press a lever to self-administer a unit dose of nicotine (30 µg/kg/injection free base; 40 µl in 1s) under a fixed-ratio schedule of reinforcement (FR) for 2-h during 10 daily sessions. During the acquisition period, the FR was set to FR1 for 5 sessions and then increased to FR2 for 5 additional sessions. Self-administration sessions were run 5–6 days/week. All self-administration sessions started with the extension of two levers and ended with their retraction after 2 h. Responding on one lever, defined as the active lever, delivered intravenous nicotine infusions, while presses on the other lever, defined as the inactive lever, had no scheduled consequence. Intravenous delivery of nicotine began immediately after completion of the active lever press and initiated a time-out (TO) period signaled by the illumination of the light cue above the lever for 20 s. Responses during the TO period were recorded but had no programmed consequence. After these 10 days of acquisition of nicotine self-administration, rats were divided in two groups and subjected to a differential dose procedure of nicotine self-administration under a FR2, a procedure previously shown to induce two between-session patterns of nicotine intake: a no-escalation (NOES) and an escalation (ES) pattern, as previously described [4]. Briefly, in the NOES group (n = 15 males and 11 females), the unit dose of nicotine remained at the initial dose of 30 µg/kg/injection for the rest of the experiment (20 additional sessions), while in the ES group (n = 14 males and 12 females) the unit dose of nicotine available progressively increased to 120 µg and then to 240 µg/kg/injection for 10 sessions at each dose. This increase in the unit dose of nicotine available was done by increasing the injection volume (from 40 to 160 and 320 µl).

#### Cue omission and nicotine omission tests

To assess the role of nicotine-associated cues and nicotine itself in the maintenance of self-administration, two test sessions were conducted: a cue omission (CueOm) and a nicotine reward omission session (NicOm), as previously described [14,15]. Following the final self-administration session, rats (NOES: n = 15 males and 11 females, ES: n = 14 males and 12 females) first underwent a CueOm test during which completion of the FR2 continued to be reinforced by an injection of nicotine (30 µg for the NOES group and 240 µg for the ES group) but without the presentation of the 20s-light cue above the lever. Conversely, in the NicOm session, completion of the FR2 was reinforced by the presentation of the cue without nicotine injections (NOES: n = 8 males and 11 females, ES: n = 8 males and 9 females). Between these two tests, rats continued to self-administer nicotine as during training for 3 sessions. Rates of operant responding during these two tests were compared to the mean of the 3 last sessions of nicotine intake escalation before the tests.

#### Nicotine-induced reinstatement of extinguished responding

After the NicOm test, rats were allowed to self-administer nicotine during three additional sessions to re-establish stable nicotine intake behavior. Next, animals (NOES: n = 15 males and 11 females, ES: n = 14 males and 11 females) underwent 5 daily extinction sessions (90 min/day) during which lever presses were recorded but no longer resulted in light-cue presentation or drug delivery. Following extinction training, rats (NOES: n = 15 males and 11 females, ES: n = 12 males and 10 females) underwent two within-session reinstatement tests (order counter-balanced between groups) with increasing doses of nicotine, as previously described [6,7]. During each reinstatement test, both levers were extended but lever pressing had no programmed consequence (no contingent drug injection or light-cue presentation). After 45 min of extinction, rats received four non-contingent doses of nicotine every 45 min (i.e., 0, 30, 60, and 120µg), each signaled by turning on the light-cue above the active lever for 20s. Increasing doses were obtained by increasing the volume of injection (0, 40, 80 and 160 µl). Drug seeking, indicated by the number lever pressing on the active lever, was measured during 45 min after each injection. In one reinstatement test, FR2 responding on the active lever had no consequence (i.e., reinstatement without response-cue contingency), while in the other, it was associated with the cue previously paired with nicotine (i.e., reinstatement with response-cue contingency).

#### Post-extinction relapse-like behavior

After the last reinstatement test, nicotine was made again available for self-administration in a subset of rats with a patent catheter (NOES: n = 15 males and 11 females, ES: n = 13 males and 9 females) during 15 consecutive sessions under the same operant conditions as before extinction. The dose of nicotine remained at 30 µg/kg/injection in the NOES, while it increased to 120 µg and then to 240 µg/kg/injection every 5 sessions in the ES group. The impact of extinction on relapse-like behavior was assessed using a within-subject comparison of the pre-versus post-extinction levels of nicotine intake.

### Drugs

(-) Nicotine hydrogen tartrate was purchased from Sigma (St. Louis, MO), dissolved in 500-ml sterile bags of 0.9% NaCl and kept at room temperature (21 ± 2°C). Nicotine doses were expressed as free base.

### Data analysis

Behavioral data were subjected to two- or three-way analyses of variance (ANOVAs) with one or two between-subject factors (Experimental groups: NOES and ES groups; sex: males and females) and one or two within-subject factors (lever: active and inactive; days: self-administration and extinction; nicotine doses: 4 doses; self-administration tests: CueOm and NicOm). All post hoc comparisons for interactions were carried out by the Newman–Keuls test. The accepted value for significance was p < 0.05. The percentages of animals that responded during reinstatement testing were analyzed using a Z-test. Trial-by-trial variability in lever press responses and nicotine injections was assessed by measuring the coefficient of variation for each variable (i.e., ratio of the standard deviation to the mean). Statistical analyses were performed using GraphPad Prism. Data graphs were created using GraphPad Prism and Inkscape.

## Results

### The initial reinforcing value of the light cue

We first assessed the initial reinforcing value of the 20-s light cue to be associated with nicotine in drug-naïve animals as a function of sex. While there was no lever preference during the initial 45-min period (sex X lever: F(1,50) = 0.004; p = 0.95, NS), female rats responded more on the light-paired lever than on the other lever during the second 45-min period (sex X lever: F(1,50) = 6.433; p = 0.01 < 0.05; active vs inactive: p < 0.001; Fig. 1b), a difference not seen in male rats (p = 0.55, NS). Importantly, both female and male rats responded equally on the inactive lever (p = 0.35; NS) and showed the same level of locomotor activity (sex effect: F(1,50) = 0.005; p = 0.95, NS; Fig. 1c). Thus, it seems that the light cue to be associated with nicotine during self-administration has some initial reinforcing value only in female rats.

### Initial acquisition of nicotine self-administration

Animals were next trained to self-administer nicotine (30 µg/kg/injection) for 10 daily sessions under a FR1 (5 sessions), then an FR2 (5 sessions) schedule of reinforcement. Both male and female rats readily acquired nicotine self-administration, demonstrated by a preference for the active lever (lever effect: F(1,100) = 116.8; p < 0.001; Fig. 2a). However, increasing response requirement to FR2 revealed sex differences in nicotine self-administration, with only females increasing accordingly their responding on the active lever (days X sex: F(9,450) = 3.57; p < 0.001). This sex difference in active responding remained stable thereafter for the rest of the FR2 sessions (sex effect: F(1,50) = 5.93; p = 0.018 < 0.05). As a result, despite an initial decrease in nicotine injections (days effect: F(9,450) = 21.93; p < 0.001; Fig. 2b), female rats self-administered more nicotine than males (sex effect: F(1,50) = 5.69; p = 0.021 < 0.05; Fig. 2c). At the end of the acquisition period, nicotine intake (i.e., average over the last 3 sessions) was higher in females than in males (F(1,50) = 7.20; p = 0.009 < 0.01; Fig. 2e). Of note, despite these differences in intake, both sexes initiated drug use equally quickly at session onset (1^st^ injection latency: F(1,50) = 0.69; p = 0.41; NS). Both sexes also increased their locomotor activity across nicotine self-administration days (days effect: F(9,450) = 12.36, p < 0.001; Fig. 2d), females being slightly more active (days X sex: F(9,450) = 2.60, p = 0.006 < 0.01). Finally, individual variation in initial responding on the light cue-paired lever (i.e., before self-administration) correlated positively with individual variation in nicotine self-administration (r = 0.40; p = 0.003 < 0.01; Fig. 2f).

**Figure 2.**
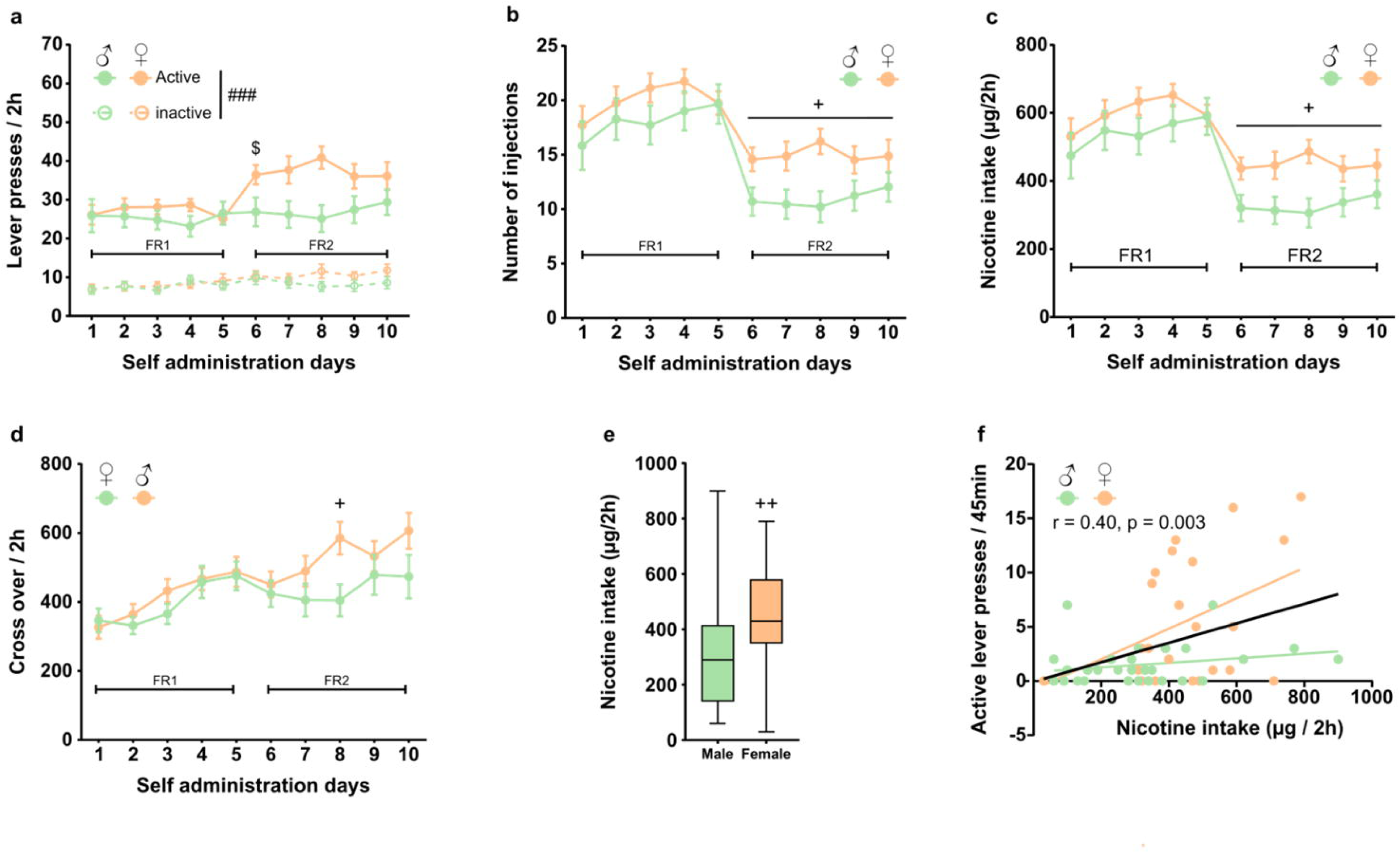
Female rats self-administer more nicotine than males during acquisition. (**a-d**) Mean (±SEM) (**a**) number of lever presses, (**b**) number of nicotine injections, (**c**) nicotine intake, and (**d**) locomotor activity during the 2 h session is plotted as a function of self-administration days for males (green, n = 29) and female (orange, n = 23) rats. (**e**) Average nicotine intake (mean ± SEM) during the last 3 days of nicotine self-administration acquisition. (**f**) Positive correlation between individual variation in initial responding on the light cue-paired lever and individual variation in nicotine self-administration. ^###^p<0.001, different from the inactive lever; ^+^p<0.05, ^++^p<0.01, different from males, ^$^p<0.05; different from session 5.

### Escalation of nicotine intake

Animals were then subjected to a recently developed differential dose procedure of nicotine self-administration [4]. Briefly, one group of rats had access to nicotine SA at the unit dose of 30µg/kg/injection (no-escalation group, NOES) during the entire procedure, while in the other group the unit dose of nicotine was progressively increased from 30µg to 120µg and to 240µg/kg/injection (escalation group, ES) to induce nicotine intake escalation. As expected, responding on the active lever decreased as the nicotine doses increased in both male ES (30µg vs 240µg: p = 0.017 < 0.05; Fig. 3a) and female ES rats (30µg vs 240µg: p< 0.001; Fig. 3b), probably through a self-titration mechanism. Responding on the inactive lever also tended to decrease but only in females (ES inactive 30µg vs 240µg, p < 0.001 and ES inactive 30µg vs 240µg, NS; for female and male respectively), resulting in a higher discrimination index in female ES than in male ES rats (120 µg active vs inactive: p <0.01, 240 µg active vs inactive: p = 0.041 <0.05 and 120 µg active vs inactive: p = 0.123, NS, 240 µg active vs inactive: p = 0.999, NS, for female and male respectively).

**Figure 3.**
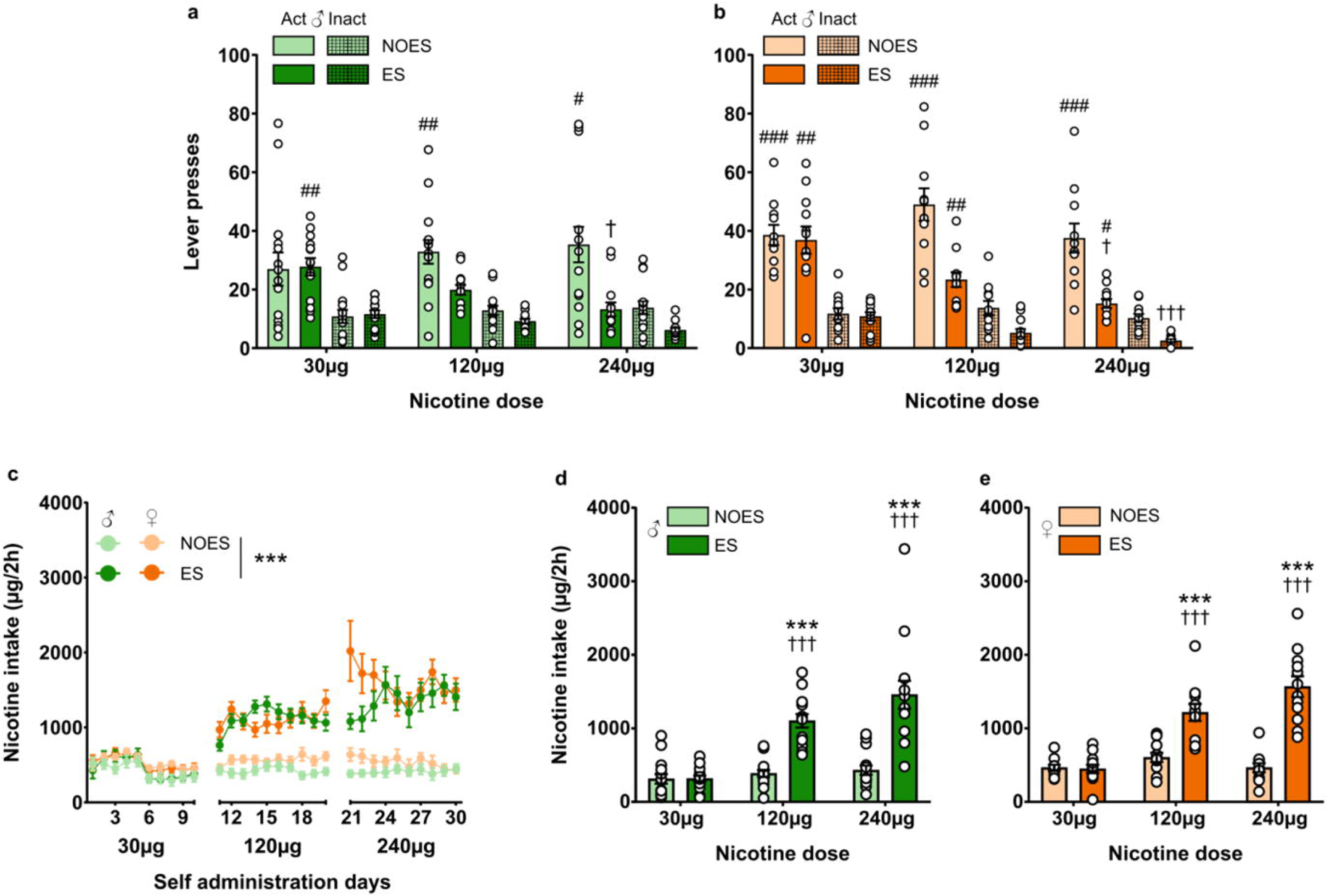
Both sexes reach similar escalated levels of nicotine intake. (**a, b**) Average lever presses (mean ± SEM) on the active (Act) inactive (Inact) lever during the last 3 days at each nicotine dose for the no-escalation group (NOES, light green and light orange) that has access to the unit dose of 30 μg/kg/injection, and rats in the escalation group (ES, dark green and dark orange) that had access to increasing doses of nicotine (30 μg, 120 μg and 240 μg/kg/injection) in males (green, NOES, n =15, ES, n = 14) and females (orange NOES, n =11, ES, n = 12). (**c**) Mean (±SEM) nicotine intake during the 2 h session is plotted as a function of self-administration days for rats in the NOES (light green and light orange) and ES rats (dark green and dark orange) (**d, e**) Average nicotine intake (mean ± SEM) during the last 3 days at each nicotine dose for the NOES and ES groups in males (**d**, green) and females (**e**, orange). ***p<0.001, different from NOES; ^#^p<0.05, ^##^p<0.01 ^###^p<0.001, different from the inactive lever; ^†^p<0.05, ^†††^p<0.001, different from 30µg.

As expected, nicotine intake remained stable in the NOES group while it increased in the ES group (group: F(1,48) = 60.750, p < 0.001; group X days F(29,1390) = 26.160, p < 0.001; Fig. 3c). However, there was no difference between males and females (sex effect: F(1,48) = 2.592, p = 0.114, NS). At the end of the dose escalation procedure, the level of nicotine intake (i.e., average over the last 3 sessions at each dose) was 3 to 4 times higher in the ES group than in the NOES group (nicotine dose X group: F(2,96) = 59.520, p < 0.001, 120µg vs 30µg ES, p < 0.001, 240µg vs 30µg ES, p < 0.01, Fig. 3d-e) with no difference between sexes (nicotine dose X group X sex: F(2,96) = 0.408, p = 0.666, NS).

### Cue omission after escalation of nicotine intake

We next assessed the contribution of nicotine and nicotine associated-cues to the maintenance of nicotine self-administration after escalation during a cue omission test (CueOm) and a nicotine reward omission test (NicOm). Cue omission (CueOm, Fig. 4a,b) decreased active lever responses in the NOES but not in the ES group (group X CueOm: F(1,48) = 6.649, p = 0.013 < 0.05; SA vs CueOm NOES, p < 0.001; SA vs CueOm ES, NS; sex X group X CueOm: F(1,48) = 0.086, p = 0.77, NS). Nicotine reward omission (NicOm, Fig. 4c,d) also decreased active responding in NOES rats (group X NicOm: F(1,32) = 17.480, p < 0.001; SA vs NicOm NOES, p = 0.019 < 0.05), but in contrast increased active responding in ES animals (SA vs NicOm ES, p = 0.034 < 0.05). These effects were observed in both males and females (sex effect: F(1,48) = 1.224, p = 0.274, NS and F(1,32) = 1.233, p = 0.275, NS, for CueOm et NicOm, respectively). Together, these results suggest that while non-escalated levels of nicotine self-administration depend on both nicotine reward and its associated cue in NOES rats, escalated levels of nicotine self-administration only depend on nicotine reward and no longer on its associated cue.

**Figure 4.**
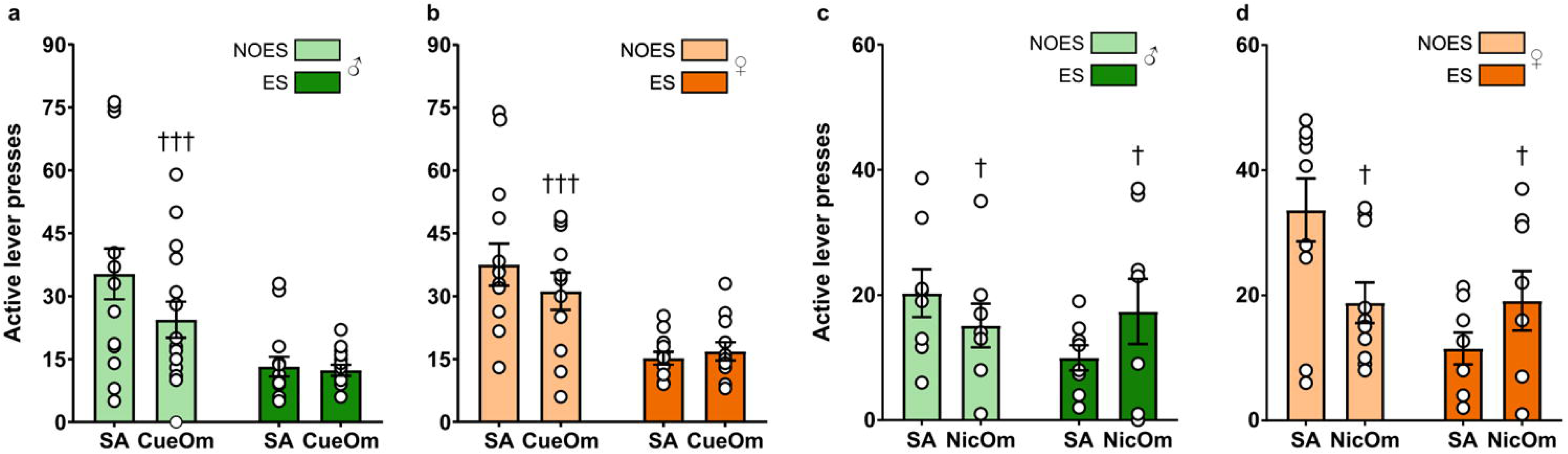
Effects of cue omission and nicotine reward omission on nicotine self-administration and escalation behaviors. Mean (±SEM) number of active lever presses during the 2h session cue omission test (**a, b**) and nicotine reward omission test (**c, d**) in NOES and ES groups in males (**a**, **c**) (green; NOES, n = 15, ES, n = 14) and females (**b, d**) (orange; NOES, n = 11, ES, n = 12). SA = mean (±SEM) of the last 3 sessions of nicotine self-administration. CueOm = Cue omission; NicOm = Nicotine reward omission. ^†^p<0.05, ^†††^p<0.001 different from SA.

### Drug seeking during extinction after escalation

Animals then underwent 5 daily sessions of extinction (90 min/day) to measure the persistence of drug-seeking behavior (Fig. 5). ES rats responded more than NOES rats on the first day of extinction (days X group: F(4, 188) = 4.86, p < 0.001; NOES vs ES, p = 0.013 < 0.05), suggesting that they were seeking the drug more. Thereafter, nicotine-seeking behavior extinguished rapidly after 3 days of extinction in the NOES group and 4 days in the ES group (days X group: F(4, 188) = 4.86, p < 0.001; NOES: Extinction session 2 vs session 3, p= 0.599 and ES: Extinction session 3 vs session 4, p = 0.614). However, there was no sex differences in the number of active lever press observed across extinction days between the two groups of animals (sex effect: F(1,47) = 0. 62, p = 0.434, NS; sex X days X group: F(4,188) = 1.22, p = 0.303, NS). Compared to active lever, there was no group (group effect: F(1,47) = 1.06, p = 0.308, NS and days X group: F(4,188) = 1.20, p = 0.311, NS) or sex difference in responding on inactive lever during extinction (sex effect: F(1,47) = 0.006, p = 0.94, NS; data not shown).

**Figure 5.**
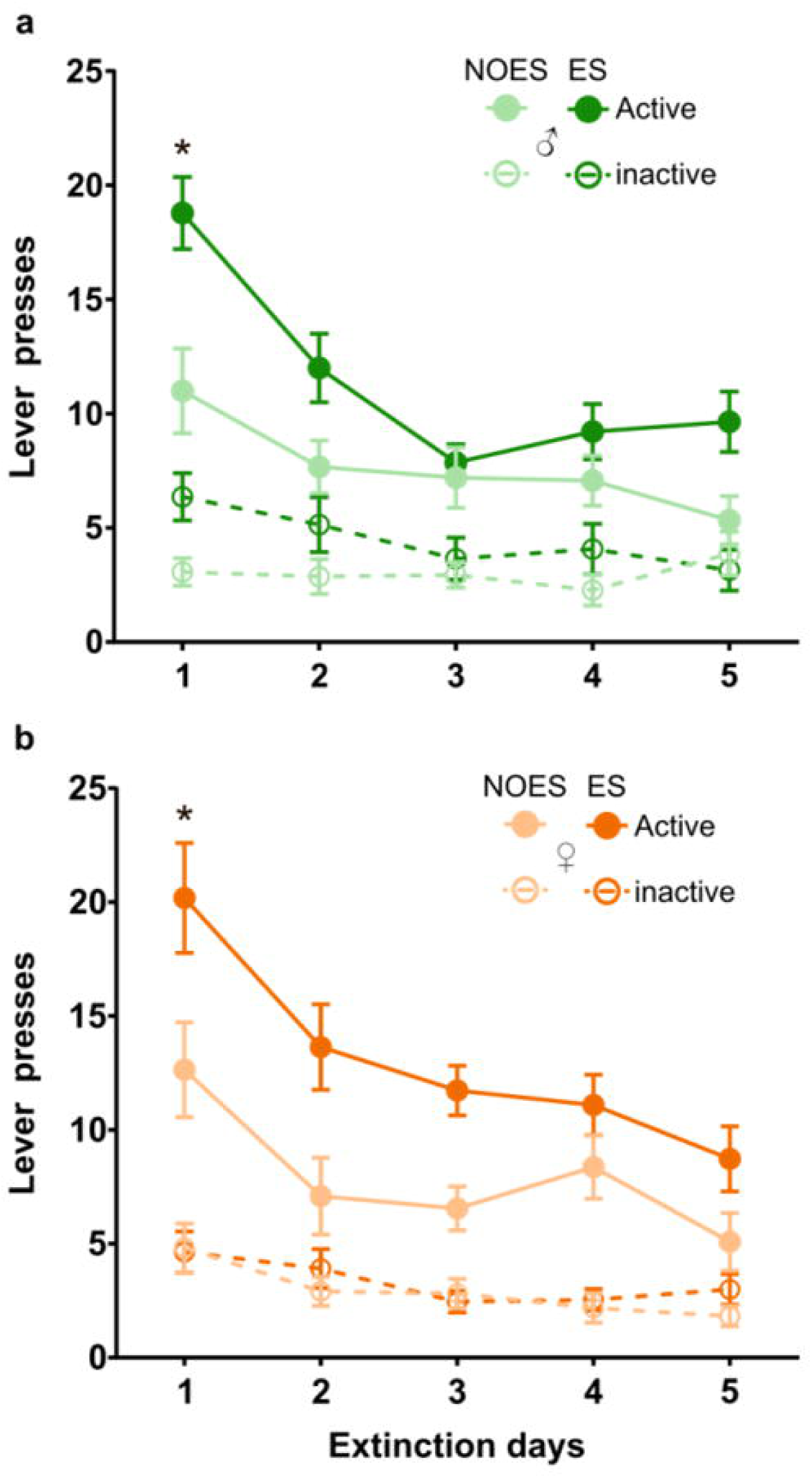
Rats show greater drug seeking during extinction after escalation. Mean (±SEM) number of active and inactive lever presses are plotted as a function of extinction days in NOES and ES groups in males (**a**) (green; NOES, n = 15, ES, n = 14) and females (**b**) (orange; NOES, n = 11, ES, n = 11). *p<0.05, different from NOES.

### Reinstatement of nicotine-seeking behavior after escalation

Nicotine-induced reinstatement was next assessed in two within-session reinstatement tests with increasing priming doses of nicotine (Fig. 6). These two reinstatement sessions were identical except that, in one of them, completion of the FR2 requirement during the 45-min inter-dose interval was associated with the 20-s light-cue presentation (i.e., response-contingent cue). In the reinstatement test without the response-contingent cue, nicotine alone only produced a small dose-dependent increase in drug-seeking (dose effect: F(3,132) = 13.52, p < 0.001; 0µg vs 120µg, p < 0.001; Fig. 6a,b) but this effect was similar in NOES and ES animals (group effect: F (1,44) = 0.15, p = 0.70, NS and group X dose: F(3,132) = 0.54, p = 0.67, NS). Interestingly, when the response-contingent cue was present, this facilitated the dose-dependent reinstatement of nicotine-seeking behavior (dose effect: F(3,132) = 8.05, p < 0.001; 0µg vs 120µg, p < 0.001; Fig. 6d,e), a facilitation mainly present in ES rats at the highest doses tested but not in NOES rats (group X dose: F(3,132) = 2.92, p = 0.036 < 0.5; ES 0µg vs ES 120µg: p < 0.01; NOES 0µg vs NOES 120µg: p = 0.25, NS). There was no sex difference in nicotine-induced reinstatement of drug-seeking behavior (sex effect: F(1,44) = 1.07, p = 0.31, NS and F(1,44) = 0.30, p = 0.59, NS, for each reinstatement test respectively). Finally, in both reinstatement tests, nicotine dose-dependently increased locomotion similarly in both groups and sexes (dose effect: F(3,132) = 37.64, p < 0.001; 30µg vs 120µg, p < 0.001; 60µg vs 120µg, p < 0.001; sex X group X dose: F(3,132) = 0.50, p = 0.68, NS; and dose effect: F(3,132) = 29.52, p < 0.001; 30µg vs 60µg, p < 0.01, 30µg vs 120µg, p < 0.001, 60µg vs 120µg, p < 0.001; sex X group X dose: F(3,132) = 0.50, p = 0.68, NS, for each reinstatement test respectively; Fig. 6c,f). These results thus suggest that the response-cue contingency during lever presses is crucial to reinstate nicotine-seeking behavior and that animals exhibited higher levels of reinstatement of nicotine-seeking behavior after escalation.

**Figure 6.**
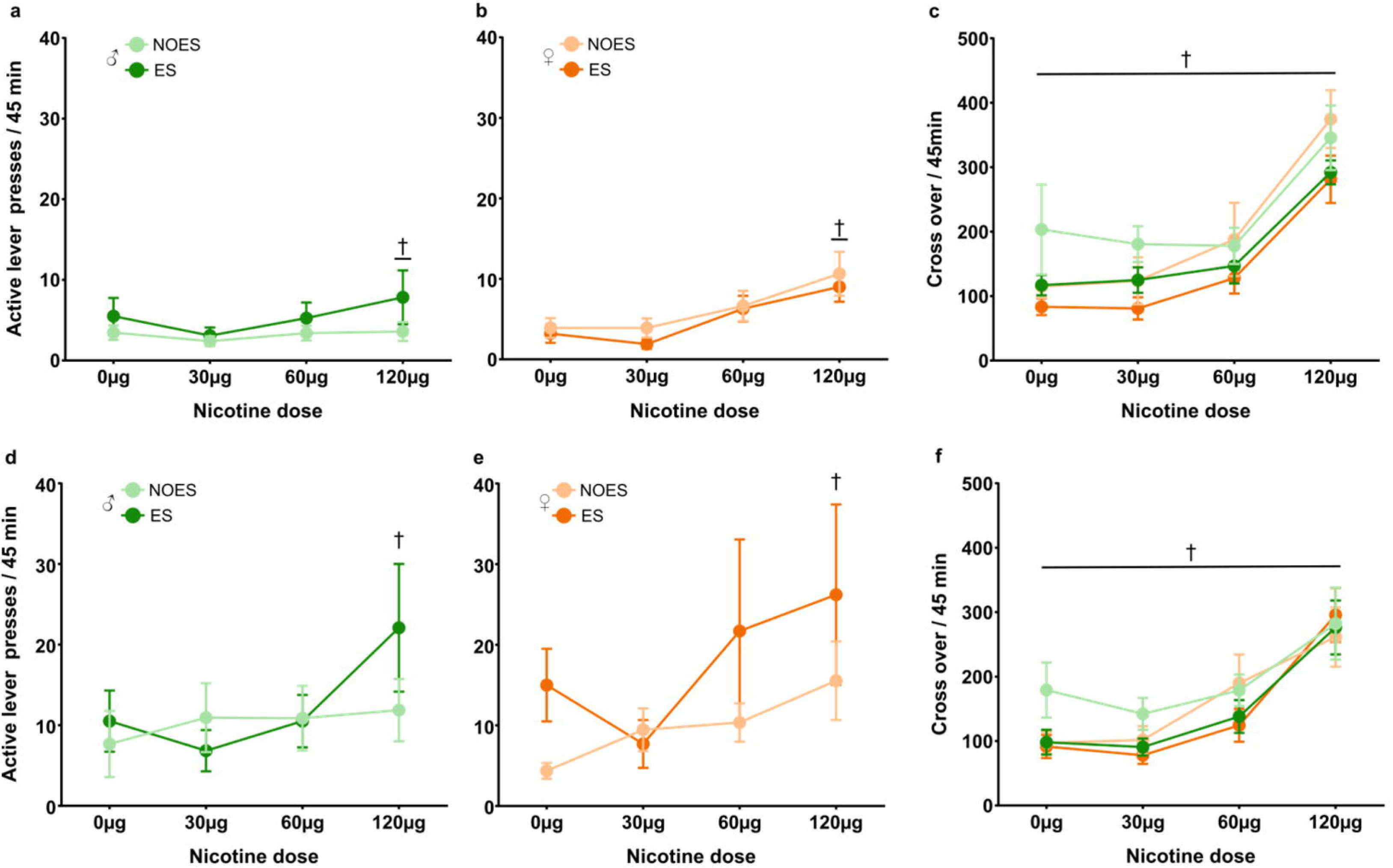
Rats show higher levels of reinstatement of nicotine-seeking behavior after escalation. Mean (±SEM) number of active lever presses during nicotine-induced reinstatement in NOES and ES groups in males (**a,d**) (green; NOES, n = 15, ES, n = 12) and females (**b,e**) (orange; NOES, n = 11, ES, n = 10). Nicotine-induced reinstatement was assessed in two within-session reinstatement tests with increasing doses of nicotine (one dose of nicotine every 45 min). In one reinstatement test, responding on the active lever had no consequence (i.e., reinstatement without response-cue contingency) (**a,b**), while in the other, it was associated with the cue previously paired with nicotine (i.e., reinstatement with response-cue contingency) (**d,e**). Mean (±SEM) number of cross over during nicotine-induced reinstatement with response-cue contingency in NOES and ES groups(**c,f**). ^†^p < 0.05, different from 0 µg.

### Re-escalation of nicotine intake after extinction

Finally, after the last reinstatement test, rats were given again access to nicotine self-administration and escalation as before extinction to measure relapse-like behavior (Fig. 7). As expected, animals in the ES group rapidly re-escalated their nicotine consumption (days X group: F(1,44) = 50.75, p < 0.001, Fig. 7a). Overall, nicotine intake at the end of resumption (i.e., average over the last 3 sessions at 240µg) was higher than nicotine intake at the end of the initial procedure in ES but not in NOES animals (group X resumption: F(1,44) = 6.800, p < 0.05; ES: SA vs Res, p < 0.001, NOES: SA vs Res, p = 0.808, NS; Fig. 7b,c) with no sex differences (sex effect: F(1,44) = 0.159, p = 0.693, NS; sex X group X resumption: F[1,44] = 0.046, p = 0.832, NS). Thus, nicotine intake re-escalated at higher levels after extinction in both male and female rats.

**Figure 7.**
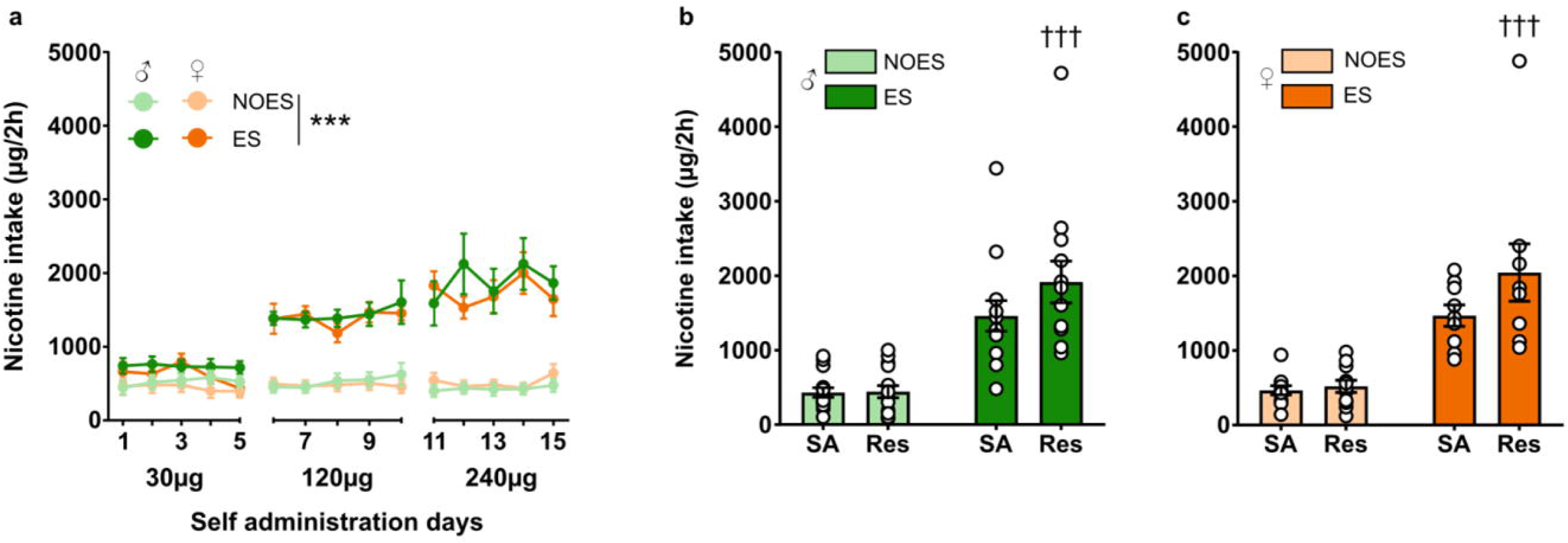
Rats re-escalate nicotine intake at higher levels after extinction. **(a)** Mean (±SEM) nicotine intake during the 2h session of nicotine self-administration after extinction is plotted as a function of days in NOES and ES groups in males (green; NOES, n = 15, ES, n = 12) and females (**b**) (orange; NOES, n = 11, ES, n =9). (**b, c**) Average nicotine intake (mean ± SEM) during the last 3 days of nicotine self-administration before extinction (SA) was compared to average nicotine intake (mean ± SEM) during the last 3 days of nicotine resumption post-extinction (Res) in NOES and ES groups in males (green; NOES, n = 15, ES, n = 12) and females (**b**) (orange; NOES, n = 11, ES, n =9). ***p<0.05, different from NOES; ^†††^p < 0.001, different from SA.

## Discussion

We first reproduced and extended our previous finding by showing that access to an increasingly large unit dose of nicotine (i.e., from 30 μg to 240 μg/kg/injection) for self-administration precipitated a robust escalation of nicotine intake in both males and female rats. However, though some sex differences existed during the acquisition of nicotine self-administration, specially under the FR2 reinforcement schedule [16–18], these differences tended to disappear during and after escalation of nicotine intake, as both male and female rats rapidly escalated their nicotine intake and reached similar post-escalation levels of intake. Lack of sex differences has also been previously observed during intake escalation of other drugs, such as, for instance, methamphetamine and heroin [19,20].

Previous studies have highlighted the critical role of nicotine-associated cues in maintaining nicotine-self-administration in male rats [10–14]. We confirmed and extended this finding to females, but only in NOES rats that did not escalate their nicotine intake. Indeed, male and female NOES rats decreased their self-administration of nicotine when the response-contingent cue was omitted. Importantly, this influence of the cue on nicotine self-administration was not observed in ES rats as their escalated levels of nicotine intake remained unchanged after cue omission. This outcome suggests that after escalation, nicotine becomes a primary source of reinforcement that no longer seems to require any facilitatory or permissive influence from external cues. This interpretation is supported by the compensatory reaction of ES rats to nicotine omission which was not observed in NOES rats. While nicotine omission resulted in a slight decrease in active lever responding in NOES rats, it led to increased responding in ES rats, probably reflecting seeking of the pharmacological effects of nicotine. Thus, it seems that after escalation of nicotine intake, escalated levels of nicotine self-administration become primarily dependent on nicotine alone and much less on its associated cues.

Consistent with the above conclusion, ES animals also responded more on the active lever than NOES animals on the 1^st^ day of extinction. Such an increase in drug-seeking has been previously observed after escalation of others drugs of abuse [5–9]. However, rather surprisingly, nicotine intake escalation had only a moderate effect on nicotine-induced reinstatement. While nicotine did not effectively prime reinstatement of nicotine seeking in absence of the response-contingent cue, even at the highest dose tested, it did in a dose-dependent manner when the cue was present, a finding that confirms previously research [21–23]. Interestingly, the latter effect was only observed in ES rats, but not in NOES rats, a difference that could not be accounted for by a difference in sensitivity to the psychomotor effects of nicotine, since nicotine increased dose-dependently locomotion equally in both groups. Overall, these findings are consistent with those observed after escalation of cocaine and heroin intake [6,7,24–26]. Again, we found no sex difference in drug-seeking or nicotine-induced reinstatement, a finding consistent with the majority of others studies on nicotine [22,27,28] but see [29]. This contrasts, however, with studies on other drugs, such as, for instance, cocaine in which females were reported to exhibit more cocaine seeking than males [30–32].

Finally, when re-exposed to the differential dose procedure, ES rats rapidly re-escalated their nicotine intake and reached higher levels of consumption than originally established before extinction, while NOES rats did not. This post extinction relapse-like behavior was independent of the sex of the animals. Such an increase in nicotine intake after an abstinence period has been previously reported in male rats with extended but not limited access to nicotine [3,29,33,34] and cocaine self-administration [35–37], suggesting that, like other drugs, a history of nicotine intake escalation increases vulnerability to relapse.

In conclusion, the present study shows that after escalation of nicotine intake, animals develop other behavioral signs of addiction, including greater drug-seeking during extinction, more intense drug-induced reinstatement, and an increased vulnerability for post-extinction re-escalation than rats with stable patterns of nicotine intake. Overall, this study also revealed no substantial sex differences in the development of these different addiction-related phenomena. It also revealed that after escalation, nicotine intake became primarily dependent on nicotine reinforcement and less so on the nicotine-paired cue. The significance and mechanisms of this loss of influence of nicotine cues on escalated levels of nicotine represent an important area of future research. Thus, overall, this study shows that most of the post-escalation behavioral changes previously seen with other drugs of abuse are generalizable to nicotine intake escalation.

## Author contributions

KG and SHA designed research and experiments; KL and LL performed behavioral experiments and associated data analysis; KL, SHA and KG wrote the paper.

